# Cytoplasmic translocation of nuclear lysine-specific demethylase-1 (LSD1/KDM1A) in human hepatoma cells is induced by its inhibitors

**DOI:** 10.1101/274050

**Authors:** Suemi Yabuta, Yoshihiro Shidoji

## Abstract

Histone-modifiable lysine-specific demethylase-1 (LSD1/KDM1A) is often upregulated in many cancers, including hepatoma, and is regarded as oncoprotein. We previously reported that the hepatoma-preventive geranylgeranoic acid (GGA) inhibits KDM1A activity at the same IC_50_ as that of the clinically used drug tranylcypromine, a verified inhibitor of KDM1A. Here, we report that these inhibitors induced cytoplasmic translocation of nuclear KDM1A in a human hepatoma-derived cell line. Immunofluorescence studies revealed cytoplasmic localization of KDM1A, 3 h after addition of GGA or tranylcypromine in HuH-7 cells. Geranylgeraniol and all-*trans* retinoic acid were both unable to induce translocation of nuclear KDM1A, whereas farnesoic acid showed the weak activity. Furthermore, GGA did not affect subcellular localization of another histone lysine-specific demethylase, KDM5A. This suggests that the inhibitor-induced translocation of nuclear KDM1A to the cytoplasm is specific for KDM1A. These data demonstrate for the first time that KDM1A inhibitors specifically induce the cytoplasmic translocation of nuclear KDM1A.

**Abbreviations:**

ATRA: all-*trans* retinoic acid
CoREST: corepressor for element 1-silencing transcription factor
DIC: differential interference contrast
FA: farnesoic acid
GGOH: geranylgeraniol
GGA: geranylgeranoic acid
LSD1/KDM1A: lysine-specific demethylase-1
pHH3: phospho-histone H3(Ser10)
TCP: *trans*-2-phenylcyclopropylamine

## Introduction

Epigenetic alterations often promote or even drive cancer development by activating and sustaining cancer-promoting gene expression.^1^ Among epigenetic changes in cancer cells, histone lysine methylation/demethylation has gained substantial attention as a possible target in therapeutic drug development.^2^ Histone lysine methylation/demethylation is extensively involved in nucleosome remodeling and gene expression. Lysine residue methylation is reversibly regulated by histone lysine methyltransferases and histone lysine demethylases (KDMs). Several KDMs have so far been identified, and lysine-specific demethylase 1 (KDM1A/LSD1) has received increasing attention since its identification in 2004.^3^ Overexpression of KDM1A is frequently observed in prostate, breast, and bladder cancers, neuroblastoma, and especially hepatoma^4^, where it correlates directly with adverse clinical outcome and inversely with differentiation.^5^ Thus, KDM1A inhibitors are of clinical interest for their anticancer role as well as their potential application in other human diseases that exhibit deregulated gene expression.^6^

Among other members of the KDM family, KDM5A/JARID1A is of particular interest because it specifically removes a methyl group from the trimethylated lysine 4 of histone H3 to produce dimethylated lysine 4. This produces a substrate for KDM1A, which is cooperatively capable of removing dimethyl and monomethyl groups from lysine 4 of histone H3.^6,7^ KDM1A and KDM5A are recruited to chromatin through the CoREST^6^ and ORC2–SUMO2^8^ complexes, respectively. Hence, these KDMs are generally recognized as nuclear-localized proteins.^9,10^

KDM1A is a flavin adenine dinucleotide-containing enzyme belonging to the amine oxidase superfamily.^11^ Structural homology between KDM1A and monoamine oxidase-B, a clinically validated pharmacological target, suggests that KDM1A is a druggable target. Indeed, screening of known monoamine oxidase inhibitors has uncovered KDM1A inhibitors that are effective at sub-millimolar concentrations, among which the best known is the clinically used antidepressant drug *trans*-2-phenylcyclopropylamine (TCP). Multiple clinical trials have begun to investigate the antitumor effect of TCP.^12^

Recently, we reported that all-*trans* geranylgeranoic acid (GGA), a natural acyclic diterpenoid found in medicinal herbs^13^, inhibits KDM1A activity at the same IC_50_ as that of TCP.^14^ GGA has been reported to induce cell death in several human hepatoma cell lines.^13,15^ Incomplete autophagic response^16^, nuclear translocation of the cytoplasmic p53^17^, and endoplasmic reticulum stress response^18^ have been explored as potentially being involved in GGA-induced cell death. Indeed, the 4,5-didehydro derivative of GGA has been repeatedly investigated and demonstrated to potentially prevent second primary hepatoma in clinical trials.^19^–^22^

To our knowledge, no studies have yet reported KDM1A inhibitor-induced changes in the subcellular distribution of the enzyme. When studying the effects of GGA on hepatoma cells, we found that GGA altered the subcellular localization of KDM1A, which could be important for both KDM1A-inhibitor screening and understanding KDM1A biology. Here, we report that some KDM1A-inhibitors can remove KDM1A from chromatin and move it to the cytoplasmic space in human hepatoma cells.

## Results

### Cytoplasmic translocation of nuclear KDM1A induced by GGA or TCP

First, nuclear localization of the chromatin protein KDM1A was confirmed in control cells by immunofluorescence. The upper row of images in Fig. 1A clearly indicates a strict colocalization of KDM1A with Hoechst 33258-stained chromatin DNA. However, 3 h after treatment of HuH-7 cells with the KDM1A inhibitors GGA or TCP, the subcellular localization of KDM1A appeared to have shifted from the nucleus to the cytoplasmic space, as shown in the lower two rows of images in Fig. 1A. Quantitative analysis of the green fluorescence intensity of KDM1A clearly shows that both inhibitors significantly affected the subcellular localization of KDM1A (Fig. 1B). This indicates that exclusion of nuclear KDM1A was induced by both KDM1A inhibitors.

**Figure 1.**
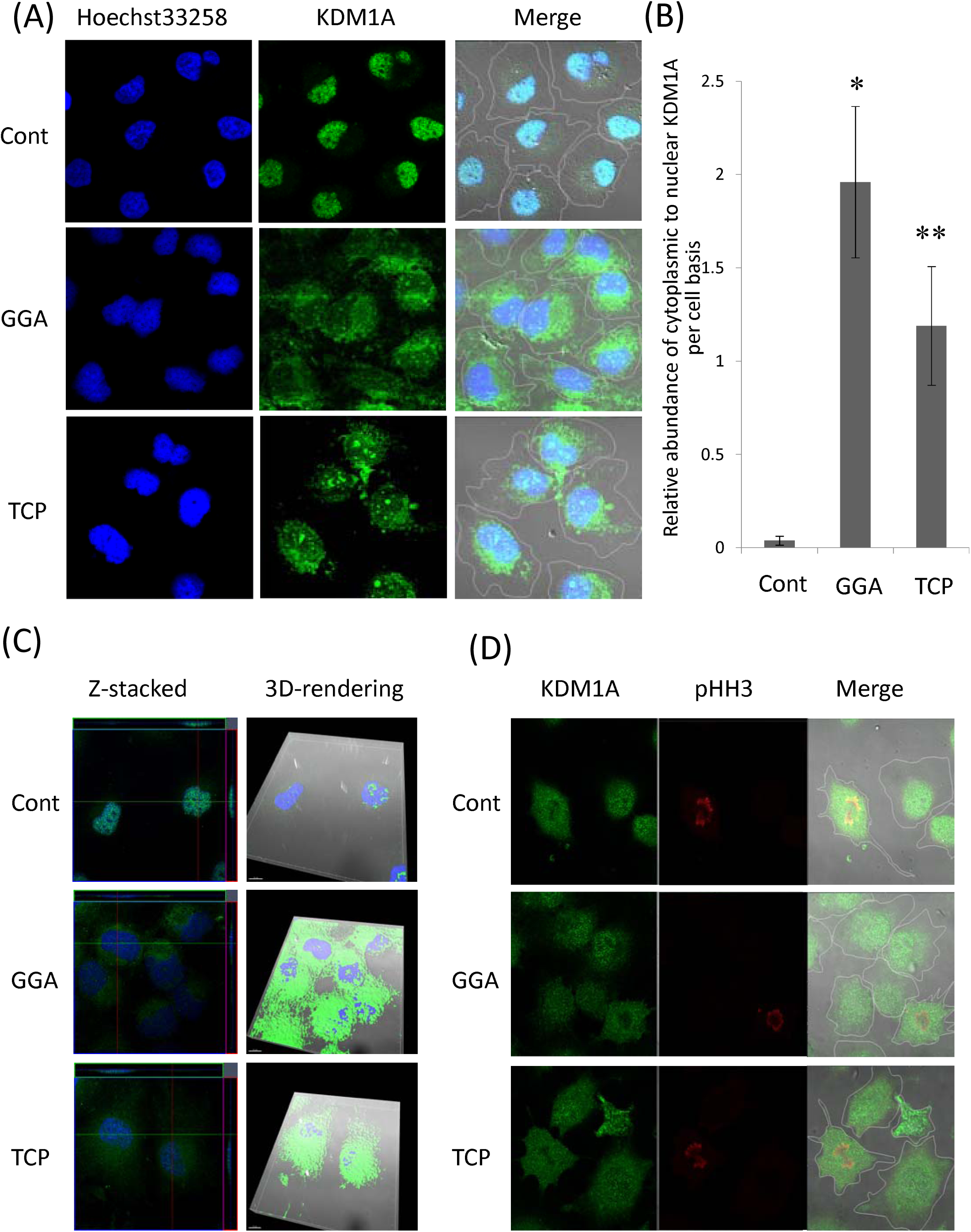
Cytoplasmic translocation of nuclear lysine-specific demethylase-1 (KDM1A) induced by either geranylgeranoic acid (GGA) or *trans*-2-phenylcyclopropylamine (TCP) treatment. HuH-7 cells were incubated for 3 h with GGA (20 μM), TCP (5 μM) or vehicle alone as a control. (A) Images were obtained of immunofluorescence staining with anti-KDM1A primary antibody and anti-rabbit IgG–Alexa488 secondary antibody (green); nuclei were counter-stained with Hoechst 33258 (blue). Merged images (Merge) were constructed using differential interference contrast (DIC) images. (B) Quantitative analysis was performed on a cellular basis with the green fluorescent images in panel (A) and relative abundance of the cytoplasmic to the nuclear fluorescence intensity was calculated. *: *P*<0.05, **: *P*<0.01 (compared with control). (C) In the merged Z-stacked images of KDM1A (green) with nucleus (blue), orthogonal hairlines show the XZ (green line and upper green box) and YZ (pink line and right pink box) planes in each panel. Z-stacked images were processed to obtain three-dimensional renderings using Imaris software. The solid view of the three-dimensional renderings allows confirmation of the cytoplasmic export of KDM1A (green) from the nucleus (blue) in GGA-and TCP-treated cells, whereas in control cells the majority of KDM1A is located in the nucleus. (D) Immunofluorescent staining of KDM1A (green) and staining of M-phase nuclei with anti-phospho-histone H3 (Ser10) (pHH3) antibody (red) in HuH-7 cells after treatment for 3 h with GGA (20 μM), TCP (5 μM) or vehicle alone as a control. White lines in the merged images mark the peripheries of cells.

The exclusion of nuclear KDM1A by treatment with its inhibitors was further confirmed by Z-axis projections (top and right parts of each image in the left column of Fig. 1C). Three-dimensional rendering of the three images using z-stacks clearly illustrates that the KDM1A protein was localized outside the nuclei 3 h after treatment with GGA or TCP, whereas most of the KDM1A protein was sequestered in the nuclei of the control cells (the right column of Fig. 1C).

Nair et al have reported that KDM1A is recruited to the chromatin of rodent embryonic stem cells in the G1/S/G2 phases and is displaced from the chromatin of M-phase cells.^23^ Therefore, we next examined whether this was the case also in human hepatoma-derived HuH-7 cells. As shown in the upper row of images in Fig. 1D, in anti-pHH3-stained M-phase cells, KDM1A protein was distributed throughout the inside of the cell, whereas in the other two cells on the right side of the M-phase cell the protein was retained in the nucleus. However, in cells treated with a KDM1A inhibitor (GGA or TCP), the protein was distributed throughout the whole cell or outside the nucleus not only in M-phase cells, but in the other remaining cells (images in lower two tiers of Fig. 1D).

### Specificity for KDM1A inhibitors and nuclear KDMs

Having identified GGA-induced translocation of nuclear KDM1A for the first time, we next considered whether this cytoplasmic translocation could be induced by treatment with a similar diterpenoid of ATRA (all-*trans* retinoic acid) that lacks KDM1A-inhibitory activity, or weak inhibitor isoprenoids of GGOH (geranylgeraniol) and FA (farnesoic acid).^14^ ATRA did not induce cytoplasmic translocation of nuclear KDM1A. GGOH and FA less-efficiently but significantly induced cytoplasmic release of nuclear KDM1A (Fig. 2A), suggesting that the translocation response of nuclear KDM1A is associated with its inhibitor activity.

**Figure 2.**
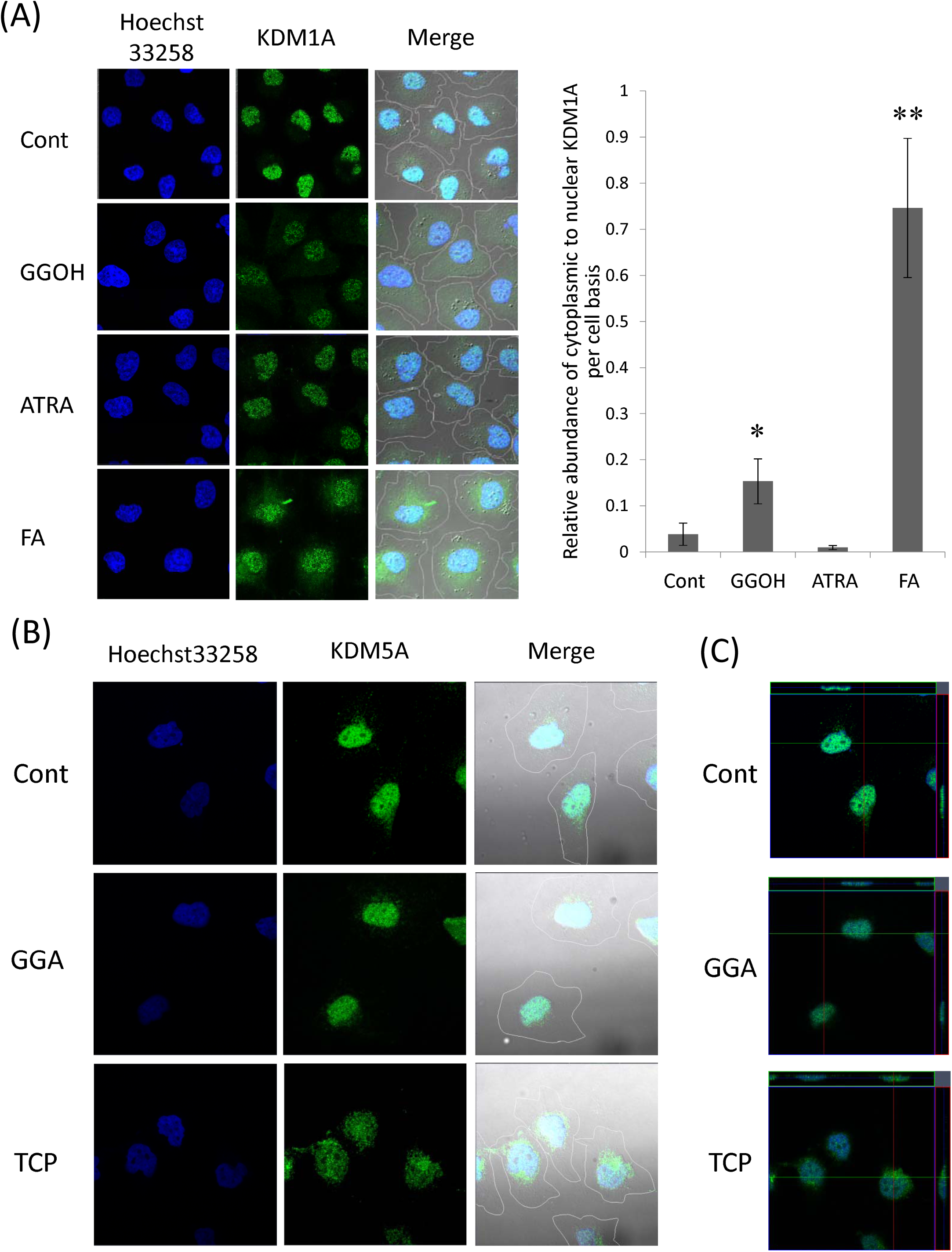
Nuclear localization of KDM1A and KDM5A after 3-h treatment with non-inhibitory isoprenoids and KDM1A-inhibitors. (A) Immunofluorescence staining of KDM1A (green) and counter-staining with Hoechst 33258 (blue) in HuH-7 cells treated for 3 h with the GGA-analogous isoprenoids geranylgeraniol (GGOH), all-*trans* retinoic acid (ATRA), and farnesoic acid (FA). Each isoprenoid was used at a final concentration of 20 μM. Quantitative analysis was performed on a cellular basis with the green fluorescent images and relative abundance of the cytoplasmic to the nuclear fluorescence intensity was calculated. *: *P*<0.05, **: *P*<0.01 (compared with control). (B) After GGA or TCP treatment, cells were immunostained for KDM5A (green), another lysine-specific demethylase, and the nuclei were counter-stained with Hoechst 33258 (blue). (C) In the merged images of KDM5A (green) and nucleus (blue) staining, orthogonal hairlines show the XZ (green line and upper green box) and YZ (pink line and right pink box) planes in each panel.

Next, we tested whether GGA or TCP could induce cytoplasmic translocation of another nuclear KDM, KDM5A, which acts cooperatively with KDM1A to remove methyl groups from histone H3 lysine 4. As shown in the upper images in Fig. 2B, KDM5A was localized only in the nuclei of control cells. Neither GGA nor TCP, inhibitors of KDM1A but not of KDM5A, induced cytoplasmic translocation of the nuclear KDM5A; this was further confirmed by Z-axis projections (top and right parts of each image in Fig. 2C).

### Release of nuclear KDM1A by GGA or TCP under cell-free conditions

We recently reported that GGA directly inhibits recombinant human KDM1A.^14^ TCP acts as an irreversible inhibitor forming a covalent adduct with the FAD cofactor of the KDM1A enzyme.^24^ In the present study, we examined the direct effect of these inhibitors on the release of KDM1A protein from the nucleus. Dose-dependent release of the protein from the nuclear fraction was observed with either GGA or TCP (Fig. 3A).

**Figure 3.**
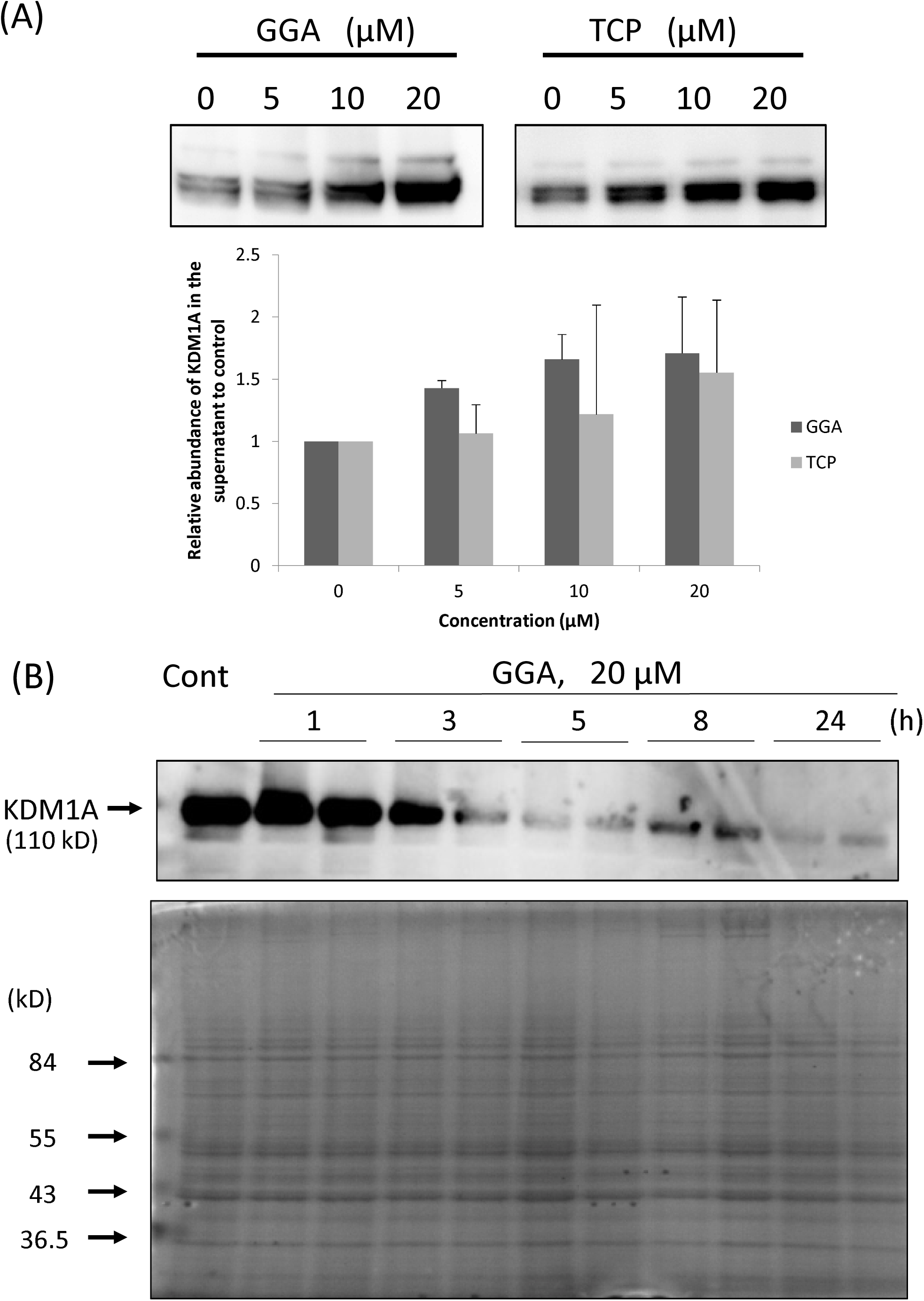
Release of nuclear KDM1A after incubation of the nuclear fraction with GGA or TCP (cell-free experiment) and downregulation of cellular KDM1A by GGA treatment (cellular experiment). (A) GGA and TCP directly released KDM1A from the nuclear fraction in a concentration-dependent manner. The nuclear fractions were treated with either GGA (5 to 20 μM) or TCP (5 to 20 μM) at 4°C for 24 h. After centrifugation, the supernatant was analyzed SDS-PAGE and western blotting. The lower bar graph shows densitometric analysis of the KDM1A bands on the blots (n = 3). (B) Western blot of KDM1A protein in whole cell lysates of HuH-7 cells treated with 20 μM GGA for the indicated times. A lower panel shows protein staining with coomassie brilliant blue.

Finally, we investigated GGA-induced changes in the cellular level of KDM1A protein to understand why KDM1A translocates out of the nucleus upon GGA treatment. Fig. 3B clearly shows that cellular levels of KDM1A decreased in a time-dependent manner after GGA treatment, which suggests that GGA may induce KDM1A protein degradation, because the *KDM1A* gene transcript level did not change during GGA treatment (unpublished data).

## Discussion

The present study demonstrates for the first time that small chemical inhibitors of KDM1A, GGA and TCP, induce cytoplasmic translocation of nuclear KDM1A protein from the nucleus in human hepatoma-derived cells. Taking into account the fact that KDM1A has been identified as a component of the corepressor for element 1-silencing transcription factor (CoREST) corepressor complex^25^ and is a recognized nuclear marker protein^26^, the present finding of KDM1A inhibitor-induced translocation of KDM1A from the nucleus to the cytoplasm may be important both for inhibitor screening and for understanding the biology of KDM1A. In this report, we propose and discuss the biological significance of the transfer of KDM1A protein from the nucleus to the cytoplasm in response to KDM1A inhibitors.

In the present study, we confirmed strict nuclear localization of KDM1A protein in control human hepatoma-derived HuH-7 cells (Fig. 1). KDM1A was discovered in 2004 as a nuclear homolog of amine oxidases that functions as a histone demethylase and transcriptional corepressor,^3^ which indicates that the enzyme originally resides in chromatin and plays a role in the repression of gene expression through epigenetic mechanisms. Here, however, we describe the striking finding that the KDM1A inhibitors GGA and TCP immediately dislodged KDM1A from the nucleus to the cytoplasm, suggesting that these inhibitors not only inhibited enzyme activity, but moved the enzyme itself to a location where epigenetic regulation was not possible.

Therefore, these two inhibitors block the function of the enzyme via a dual mechanism.The question then arises as to what types of compounds can induce this transfer of the KDM1A enzyme from the nucleus to the cytoplasm. We tested the ability of the GGA-analogous compounds GGOH (an alcoholic derivative of GGA), ATRA (a monocyclic and conjugated derivative of GGA) and FA (an acyclic sesquiterpenoid acid, whereas GGA is an acyclic diterpenoid acid) to induce the cytoplasmic translocation of nuclear KDM1A. ATRA, which exhibit no KDM1A-inhibitory activity, failed to induce translocation of the enzyme and both GGOH and FA with less inhibitory activity^14^ slightly but significantly translocated the enzyme (Fig. 2), suggesting that the activity of the inhibitors may be associated with the dislodging activity. Hence, one can reasonably speculate that an assay for drug-induced translocation of nuclear KDM1A could be developed to screen for KDM1A inhibitors without the need for measurement of the enzyme activity.

A second question is whether these enzyme inhibitors dislodge only KDM1A from the chromatin. We investigated another lysine-specific demethylase, KDM5A, which plays a cooperative role in demethylation of histone H3 lysine 4 in conjunction with KDM1A.^27^ We confirmed that KDM5A localized to the nucleus. Neither GGA nor TCP induced translocation of the nuclear KDM5A to the cytoplasm under the same conditions as those under which these inhibitors dislodged nuclear KDM1A. This indicates that the inhibitors’ effect on subcellular localization is specific for KDM1A. However, it currently remains unclear whether other components, such as HDAC1/2, may be released from the CoREST complex by GGA or TCP treatment.

Recruitment of KDM1A to chromatin has been extensively and thoroughly explored^28–32^, but little research has been conducted on the unloading of KDM1A from chromatin. Particularly relevant among the previous studies on the subcellular distribution of KDM1A is a report by Nair et al, which describes cell cycle-dependent association and dissociation of KDM1A with chromatin.^23^ Specifically, KDM1A is recruited to the chromatin of cells in the G_1_/S/G_2_ phases and is displaced from the chromatin of M-phase cells. We confirmed that in M-phase cells stained with pHH3 antibody, KDM1A protein was distributed throughout the cell (Fig. 1D), whereas in the other non-M-phase cells KDM1A protein remained in the nucleus. However, in cells treated with GGA or TCP, KDM1A was distributed throughout the whole cell interior as in control M-phase cells, regardless of cell cycle stage. We can therefore conclude that GGA and TCP did not increase the number of cells in M-phase, and did act on cells in the G_1_/S/G_2_ phases to induce translocation of nuclear KDM1A to the cytoplasm. Taking these findings together with the fact that anti-pHH3 antibody recognizes phosphorylated Ser10 on histone H3 and that introduction of a negatively charged phosphate group on Ser10 completely abolishes KDM1A activity^33^, one can reasonably speculate that a KDM1A inhibitor may unload the enzyme from chromatin.

Finally, we should discuss how GGA and TCP specifically dislodged KDM1A from the nucleus. We speculated that direct binding of GGA or TCP to nuclear KDM1A might cause release of the enzyme from the chromatin. By analyzing the nuclear fraction containing KDM1A, we demonstrated that GGA-and TCP-mediated release of KDM1A from the nuclear fraction to the supernatant was dose-dependent (Fig. 3A). Both GGA^14^ and TCP^24^ directly inhibit recombinant KDM1A enzyme activity and are thought to bind to the substrate-binding site of the enzyme. The structures of free and CoREST-bound KDM1A are virtually identical, with only a small difference in the orientations of two α-helices of the tower domain relative to the amine oxidase domain of KDM1A.^34^ In this context, it is difficult to speculate as to whether direct binding of GGA or TCP to the substrate-binding site in the amine oxidase domain of KDM1A would affect interaction between the enzyme and the CoREST protein. Other inhibitor-induced conformational changes of KDM1A may be involved in its unloading from the nuclear fraction in the cell-free experiment. We should also consider the metabolic fate of the KDM1A exported in response to treatment with its inhibitor. Fig. 3B clearly shows that GGA-induced downregulation of the cellular KDM1A level was time-dependent. This may potentially be explained by polyubiquitination and efficient degradation of the dislodged KDM1A from the chromatin by the proteasome system.^35^

The present study clearly demonstrates that the KDM1A inhibitors GGA and TCP induce cytoplasmic translocation of nuclear KDM1A. This biological process has potential to be useful for screening promising cancer-preventive epigenetic therapeutic agents targeting KDM1A, and further work to this end is warranted.

## Material and Methods

### Materials

GGA and FA were generous gifts from Kuraray (Okayama, Japan). GGOH and TCP hydrochloride were purchased from Sigma Aldrich (St. Louis, MO, USA). ATRA was obtained from Wako Pure Chemicals (Osaka, Japan).

### Cell culture and treatment

Human hepatoma-derived HuH-7 cells were seeded at 1.5 × 10^4^ cells/cm^3^ and cultured in Dulbecco’s modified Eagle medium (DMEM; 4500 mg/L glucose, Wako Pure Chemicals) containing 5% heat-inactivated fetal bovine serum (FBS; Thermo Scientific Hyclone, Yokohama, Japan) for 2 d. Thereafter, the medium was replaced with FBS-free DMEM one day before introducing of relevant test compounds at the required concentration. Ethanol (0.1%, v/v) was used as a vehicle control.

### Immunofluorescence

After drug treatment, cells cultured on glass inserts in a 24-well plate were rinsed with PBS (-), the cells were incubated at 4°C overnight with the primary antibodies. The primary antibodies used were rabbit polyclonal anti-LSD1 (or KDM1A; Cell Signaling Technology, Boston, MA, USA) and rabbit monoclonal anti-JARID1A antibody (KDM5A; D28B10, Cell Signaling Technology). After being washed with PBS(-), the cells were incubated at room temperature for 1 h with Alexa Fluor 488-labeled goat anti-rabbit IgG antibody (Invitrogen, Molecular Probes, Tokyo, Japan) and/or anti-phospho-histone H3(Ser10) (pHH3)–Alexa Fluor^^®^^ 647 conjugate (Cell Signaling Technology) and the nuclei were counter-stained with Hoechst 33258 (Invitrogen). After being washed with PBS(-), the cells were mounted in PermaFluor (Beckman Coulter, Brea, CA, USA), covered with a glass slide, and observed under an LSM700 2Ch URGB confocal laser-scanning fluorescence microscope equipped with an Axio Observer Z1 Bio (Carl Zeiss, Göttingen, Germany). Microscopic images were acquired and quantitatively analyzed using Zen 2010B SP1 software (LSM700 version 6.0, Carl Zeiss). A Z-stack procedure was employed, and the acquired images were processed with Zen 2010B SP1 and three-dimensional rendering was performed with Imaris x64 version 7.1.0 (Bitplane Scientific Software, Zurich, Switzerland).

### Cell-free analysis of drug-induced release of nuclear KDM1A

Nuclear fractions of HuH-7 cells were prepared for cell-free analysis of inhibitor-induced translocation of nuclear KDM1A using a CelLytic nuclear kit (Sigma Aldrich). The nuclear pellets were gently suspended in PBS-T (PBS containing 0.1% Triton-X 100), then GGA or TCP (final concentrations 0–20 μM) was added to each nuclear suspension (60 μg protein/tube, proteins were quantified by Bradford assay, Bio-Rad, Hercules, CA, USA), and samples were gently vortexed and incubated overnight at 4°C. The supernatants were centrifuged at 1000 × g for 10 min, separated by SDS-PAGE, and analyzed by western blotting with anti-KDM1A antibody as described below.

### Western blotting

The eluted proteins in equal volume were separated by SDS-PAGE and transferred to semi-dry blotted polyvinylidene fluoride membranes (Bio-Rad). The membranes were probed with a rabbit polyclonal antibody against KDM1A (Cell Signaling Technology), and then with horseradish peroxidase (HRP)-labeled secondary antibody (GE Healthcare, Tokyo, Japan). KDM1A bands were detected with Immobilon Western Chemiluminescent HRP substrate (Merck Millipore Japan) using an Image Quant LAS 4000 (GE Healthcare).

### Statistical analysis

Differences between two groups were assessed by Student *t*-test and were considered statistically significant if *P* value was <0.05.

